# QuartPlotR: A quarternary phase diagram tool

**DOI:** 10.1101/2024.03.22.586216

**Authors:** Alaguraj Veluchamy, Chris Bowler

## Abstract

**Motivation:** Large scale studies involving exploratory data analysis and important key discoveries require platform that provides comprehensive visualization. Density distribution analysis across multiple datasets is intuitive and summarization, visualization could reveal several biological information. Integration and visualization of sequence and annotation features in the context of composition of genomic mutation, microbiota, population are significantly challenging.

**Results:** We propose a simple, novel strategy of visualization of multidimensional datasets involving multiple layers of data distribution which are interconnected. Also, we have implemented this phase diagram in an easy-to-use tool QuartPlotR, a resource for plotting charts from different genomic datasets. A generic data access and plotting framework has been designed and this is implemented as an R package.

**Availability:** https://github.com/AlagurajVeluchamy/QuartPlotR.

**Supplementary information:** Supplementary data are available at *Bioinformatics* online.

## 1 Introduction

1. About complexity of data: Increasing number of Big Data, data types and their complexity challenges computational biologists to create novel methods of both analysis and visualization. Data complexity is intrinsic to biological systems with multiple layers of interconnected data. For example, comparison of multiple quantitative data involves visualization by displaying relative abundance using bar charts or area plots or pie-charts.
2. Visualization problem: Understanding of these multiple datasets from multiple different samples require both global and in-depth view. Insights-driven data visualization simplifies decision making for biologists by increasing degrees of abstraction and reducing the complexity parameter.
3. Sample data visualization: DataVis (data visualization) is a subdiscipline of bioinformatics and provide scientific communities with plethora of choices of tools for dissemination of science visually (O’Donoghue, 2021). Additionally, Big Data such as metagenomics data encompass attributes which are both quantitative and categorical in nature. Tools such as Krona provide charts which provides interactive visualization of multiple locations (Ondov *et al*., 2011). Genetic data visualization tools for health and diseases usually are landscape based on a particular region (Jia *et al*., 2020). Shortcomings from Linear layout in displaying large volume of genomic data were mitigated in CIRCOS in the form of circular layout (Krzywinski *et al*., 2009). Spatial visualization of scRNA data such as t-SNE from multiple cancer types involves classification and comparison of commonalities and differences of individual cells in a two-dimensional map (Li *et al*., 2017).
4. What is lacking: However, these tools lack the features for displaying secondary variables that include comparing relationships between taxonomic and functional variation of multiple samples such as location of samples in metagenomic data, mutations in population with stroke or cancer.
5. About Quarternary plot: Here, we present a novel method of visualization data which are of comparative level of complexity. We demonstrated the usability of this tool using both metagenome and mutation data. Our method, Quarternary plot is a graphical representation that provides comparative visualization of both taxonomic and functional hierarchies. This plotting system is coordinate based and presented in a spatial layout as opposed to the linear and circular layouts discussed earlier. Edges of the quadrilateral is the sample. This plotter is easy to use on multiple platforms.

## 2 Methods

Phase diagram: Data points are represented as a spatial diagram in a polygon (quadrilateral). Four samples are defined as vertex of this quadrilateral. Initially, the quadrilateral is split into four triangles. For each data point, the data is represented as a centroid point, calculated from three sides of a triangle.

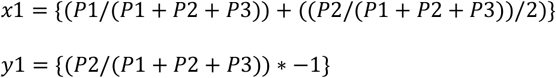

where G(x1,y1) is the centroid of the triangle. P1, P2, P3 are values of individual samples. x1, y1 are co-ordinates for each triangle. Similarly, x2, y2, x3, y3, x4, y4 are calculated for four triangles (upper, lower, left and right triangles) in a quadrilateral.

Hierarchical data abstraction: Four layers of abstraction can be visualized:

1. Data points are distributed based on the above barometric calculation.
2. Data points could be displayed as points or differential circles.
3. Size of the circle is based on the quantitation.
4. Pie chart for numerical proportion of data.

Implementation in R: Lines connecting vertex of the polygon is drawn. A grid is drawn followed by labeling of the pie-chart is done.

Data sources: Metagenome and mutation data: Metagenome data from four locations were taken and the data for tag assignment (MetaG) and metatranscriptome (MetaT) are used to plot (de Vargas *et al*., 2015).

## 3 Results

QuartPlotR accurately describes a four-sided dataset. This tool enables scientists to have an intuitive pattern from large list of genes. Visualization properties such as data points could be either dot or pie-charts. The image attributes include:

- Four corners are represented as a vertex in the diamond shaped quadrilateral.
- Data points can be displayed either as Pie chart or dot plot.
- Size of pie represents one quantitative selection.
- Data points which are located towards the corner are barometric in nature.
- When all four parts are equally shared, the points are in the middle.
- The points will be on the line, if data points shared among two only vertex only.

### 3.1 Data Structure

QuartPlotR requires a minimum of four column (tab delimited) text file (TXT). Data structure should be hierarchically defined.

- Each column could be either category or numerical.
- Minimum numerical value should be presented in all samples.

### 3.2 Illustration using metagenomics and mutation data

Metagenomics data collected from (de Vargas *et al*., 2015). As described in ocean plankton distribution, the station22, has a very high proportion of plankton which is in the range of 20-80 micron (Fig.1).

**Fig. 1.**
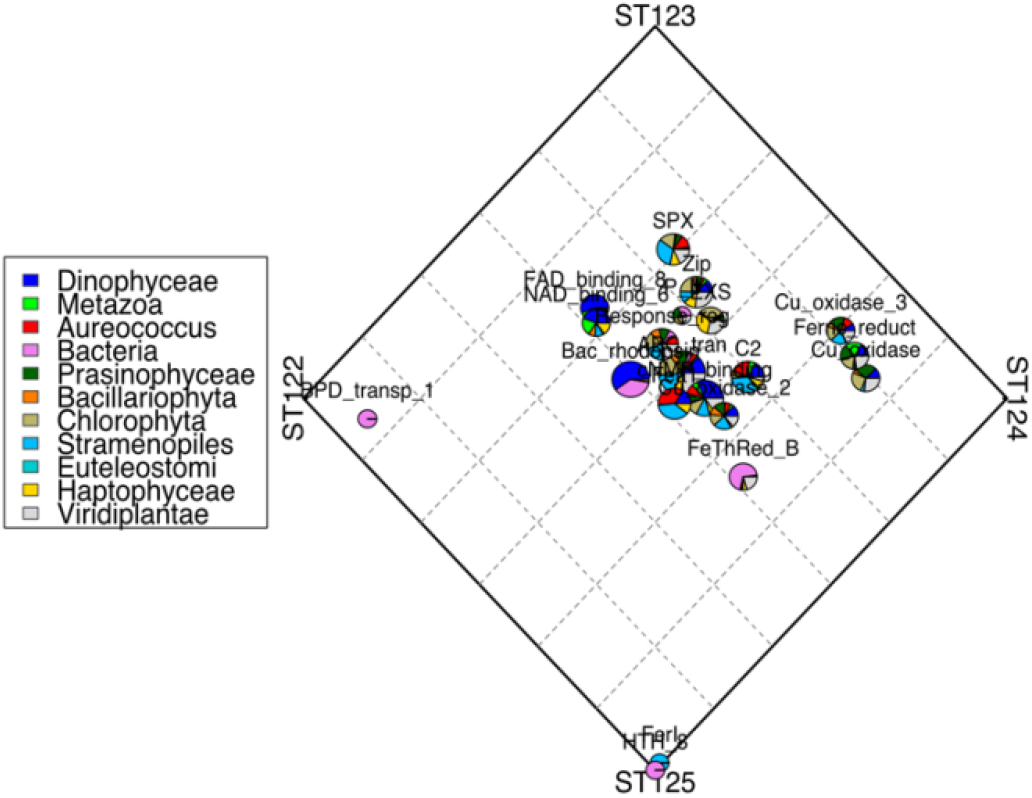
Quarternary diagram for a metatranscriptome. Using QuartPlotR to compare four metatranscriptomes tag assignment. The scale in the four quadrant refers to different station of sampling. ST122, ST123, ST124, ST125 refers to four different sampling locations.

### 3.3 Advantages and limitations

Most combination of categorical andquantitative data can be optimally visualized. Limitations include

- Comparative visualization is limited to 4 sides only.
- Large number of overlapping data points could clutter pie charts, user need to adapt dot plot in that case.
- Atleast data from two sides should be shared, otherwise datafondsThe

## References

Jia, W. et al. (2020) Oviz-Bio: a web-based platform for interactive cancer genomics data visualization. Nucleic Acids Res., 48, W415–W426.

Krzywinski, M. et al. (2009) Circos: An information aesthetic for comparative genomics. Genome Res., 19, 1639–1645.

Li, W. et al. (2017) Application of t-SNE to human genetic data. J. Bioinform. Comput. Biol., 15, 1750017.

O’Donoghue, S.I. (2021) Grand Challenges in Bioinformatics Data Visualization. Front. Bioinforma., 1, 13.

Ondov, B.D. et al. (2011) Interactive metagenomic visualization in a Web browser. BMC Bioinformatics, 12, 385.

de Vargas, C. et al. (2015) Ocean plankton. Eukaryotic plankton diversity in the sunlit ocean. Science, 348, 1261605.

